# Dissociation Between Genetic Risk and Transcriptional Output in Schizophrenia: A Cross-Tissue Meta-Analysis of *CSMD1* and *CSMD2* Expression

**DOI:** 10.64898/2026.03.18.709827

**Authors:** Mohamed-ElMehdi Boughanmi, Marion Leboyer, Caroline Demily, Romain Rey

## Abstract

**Background:** Schizophrenia is a neurodevelopmental disorder shaped by immune-related mechanisms, particularly dysregulated complement-mediated synaptic pruning. Genome-wide association studies have identified *CSMD1* as a major schizophrenia risk gene, an association robustly replicated across populations of diverse ancestries. As a complement regulator, *CSMD1* further links genetic vulnerability to synaptic refinement processes. However, the transcriptional status of *CSMD1* and its homolog *CSMD2* in individuals with schizophrenia (SZ individuals) remains poorly characterized. We conducted a meta-analysis of gene-expression datasets to determine whether *CSMD1* and *CSMD2* are differentially expressed in brain and peripheral tissues, and to assess the concordance between central and peripheral transcriptional signals.

**Methods:** Transcriptional data were obtained from gene expression omnibus. Random-effects meta-analyses were performed on *CSMD1* and *CSMD2* expression data from 854 postmortem brain samples derived from 348 SZ individuals and 346 healthy controls (HC), and 295 peripheral blood samples from 162 SZ individuals and 133 HC. Sex-stratified analyses and meta-regressions evaluated potential moderators.

**Results:** In brain tissues, *CSMD2* expression was significantly increased in SZ individuals vs. HC (SMD: 0.22 [0.05; 0.39], adj-p=0.026), whereas *CSMD1* showed no differential expression. The female-only meta-analysis revealed nominal *CSMD2* overexpression (p=0.037) in brain tissues, not surviving correction. No significant transcriptional differences were detected in peripheral blood.

**Conclusion:** In schizophrenia, our findings point to a dissociation between genetic vulnerability and transcriptional activity within the *CSMD* gene family. Schizophrenia is associated with selective brain *CSMD2* overexpression, contrasting with unchanged *CSMD1* transcription and absent peripheral blood alterations. These findings support complement-related dysregulation as a central pathway in schizophrenia.

## INTRODUCTION

Schizophrenia (SZ) is a chronic and disabling neurodevelopmental disorder characterized by positive and negative symptoms, disorganization, cognitive impairments, and social dysfunction, with a lifetime prevalence of approximately 1% and a high heritability estimated at around 80% [1, 2]. SZ highly contributes to the public health burden worldwide and is responsible for a dramatic increase in mortality due its strong association with somatic diseases, especially those related to inflammatory processes including metabolic, cardiovascular disorders and cancers [3]. Accordingly, SZ is now considered as a whole body disorder in which pathological processes take place not only at the cerebral level but also in peripheral tissues [4]. Although SZ pathogenesis is partly understood, there is strong evidence that it is a complex disorder resulting from genetic and environmental interplays [5].

Accumulating results suggest that the effects of SZ genetic risk factors manifest as postnatal neurodevelopmental alterations [6]. One of the core processes of postnatal brain maturation consists in the selective removal of unnecessary synapses while functionally important ones are strengthened, this so-called *synaptic pruning* is necessary for the setting-up of appropriate neural circuits. Increasing evidence support that a disruption of synaptic pruning during adolescence pave the way towards SZ disorders [7, 8]. Indeed, fewer synapses and a decreased number of dendritic spines are the most replicated structural findings in the postmortem brain of individuals with schizophrenia (SZ individuals) compared to heathy controls (HC) [9, 10]. Such a reduction in synapses may be morphologically reflected by decreased cortical thickness [11] which is typically observed in SZ individuals [12], and throughout different stages of SZ disorder i.e. in clinical high-risk subjects [13] and first-episode patients [14].

Although the exact mechanisms involved in the synaptic pruning remain partly understood, this process appears to rely on the joint action of the Complement system and microglia [15]. The Complement system consists of at least 30 circulating and cell-surface proteins that interact with each other leading to the elimination of pathogens while protecting the self from subsequent damage [16]. In addition to its major role in the immune surveillance, the Complement system has been implicated in the development of the central nervous system [15, 17, 18]. Especially, the Complement classical pathway is critically involved in synaptic pruning since its initiation leads to Complement component 4 (C4) cleavage which in turn promotes C3 activation and deposition on unnecessary synapses thereby tagged for elimination by microglia [7, 18, 19].

From a genomic perspective, accumulating evidence supports alterations of the Complement system in the brain of SZ individuals. The strongest genetic association with SZ risk identified by Genome Wide Association Studies (GWAS) involves variation in the major histocompatibility complex (MHC) locus [20], partly driven by structurally distinct alleles of *C4* genes [19]. More precisely, each of the four common *C4* alleles is associated with SZ risk in proportion to its tendency to induce greater expression of *C4A* in the brain [19, 21, 22]. Consistently, an overexpression of *C4* genes (*C4A* and/or *C4B*) has been observed in the total cerebral cortex, as well as in various brain regions of SZ individuals as compared to HC [19, 21, 23, 24]. Moreover, *C4A* gene copy number was recently correlated with neuropil contraction in SZ individuals [25]. Altogether, these data suggest that SZ pathogenesis is associated with an excessive Complement system activity (CSA) in brain tissues [26].

Beyond the MHC region, GWAS have identified an association between the rs10503253 variant in the complement-related gene *CSMD1* and schizophrenia risk [20], an observation subsequently replicated in candidate-gene studies across populations of diverse ancestral backgrounds [27–29]. In addition, a candidate-gene study reported associations between SZ risk and multiple variants in *CSMD2* of which rs911213 reached a statistical significance comparable to that of a genome wide threshold [29]. *CSMD1* and *CSMD2* are homologous genes encoding complement control proteins (CCP) that tightly regulate the complement system, thereby protecting host tissues from excessive or inappropriate activation and restricting complement deposition on self-surfaces [30, 31]. While CSMD1 protein inhibits classical complement pathway leading to the degradation of C3 and C4 components and regulates complement-mediated synapse elimination in the brain during development [32], CSMD2 exact role in the complement regulation is yet to be described [31].

In SZ, while the hypothesis of an increased brain CSA is supported by *C4* overexpression in various brain regions of SZ individuals [19, 21, 23, 24], studies investigating *CSMD1* and *CSMD2* expression are lacking. To date, only 3 studies explored *CSMD1* expression in the peripheral blood of small-sized samples and reported an underexpression in individuals with SZ or FEP vs. HC [33–35]. To our knowledge, no study has yet investigated *CSMD2* expression or the brain expression of *CSMD1* in SZ. Further research is therefore needed to determine whether *CSMD1* and *CSMD2* expression is altered in the brains of SZ individuals and to assess their expression in peripheral blood, to replicate previous findings. Peripheral blood is indeed considered as readily accessible surrogate system to study the molecular mechanisms involved in SZ pathogenesis [36].

Using independent gene expression datasets, we first realized a meta-analysis of the transcriptional expression of *CSMD1* and *CSMD2* in the brain tissues of SZ individuals as compared to HC. This meta-analytic approach was then performed in the peripheral blood compartment of SZ individuals *vs*. HC. Thirdly, to evaluate the neurobiological relevance of the blood results, we compared them to those observed at the brain level.

## METHODS

### 1. Search and inclusion criteria of gene expression datasets

Publicly available gene expression datasets were identified in the NCBI Gene Expression Omnibus (GEO) database (http://www.ncbi.nlm.nih.gov/geo/) until May 1st, 2023, using the following search terms: ((“schizophrenia” [MeSH Terms] OR schizophrenia [All Fields]) AND “Homo sapiens” [Filter] AND (“gse” [Filter] AND “Expression profiling by array” [Filter]). Included datasets fulfilled the following criteria: (i) derived from original studies, (ii) containing two groups (SZ individuals and HC), and (iv) obtained from brain or blood samples. Datasets investigating only non-coding RNA or using genetically reprogrammed / cultured cells were excluded.

### 2. Ethical statement

The datasets included in the present study were obtained from previous projects, all of which in accordance with ethical standards [37–50].

### 4. Data extraction

For each included dataset, sample type, platform, number of cases and controls, demographic characteristics of the subjects (sex, age, treatment), quality data of the samples (postmortem interval [PMI], pH, RNA integrity number [RIN]), and preprocessed expression level of each candidate gene were extracted using shinyGEO [51]. ShinyGEO is an application implemented using R [https://www.r-project.org/], shiny [http://shiny.rstudio.com/] and GEOquery package [52]. Common gene symbols were used to download corresponding probes.

### 5. Statistical analyses

We performed primary sex-combined meta-analyses, followed by secondary sex-stratified meta-analyses in males and females separately. Meta-analyses were carried out using the meta package on R [53]. The effect size (Hedges’s g), defined as the standardized mean difference (SMD) between gene expression levels in SZ individuals and HC samples, was calculated separately for each dataset using the metacont() function from the R meta package, along with 95% confidence intervals (CI). All analyses were conducted using a random-effects model, to account for potential heterogeneity between studies [54]. To account for multiple comparisons across candidate genes, we applied the Benjamini–Hochberg false discovery rate (FDR) correction to all p-values obtained from the meta-analyses. Adjusted p-values (adj-p) <0.05 were considered statistically significant.

Cochran’s Q-test and the I-squared (I2) statistic were computed within the same modeling framework using the meta package, providing consistent estimates of between-study heterogeneity. Q follows a chi-squared distribution with k–1 degrees of freedom; high Q-values indicate that the observed variability among studies exceeds what would be expected by chance. The I2 statistic represents the percentage of total variation across studies due to heterogeneity rather than sampling error, ranging from 0% (no heterogeneity) to 100% (considerable heterogeneity) [55]. Potential publication bias was assessed by visual inspection of funnel plots and Egger’s regression test, with a one-sided *p*-value ≤0.10 considered indicative of asymmetry [56]. When bias was detected, the trim-and-fill method was applied to estimate and adjust for the potential influence of missing studies.

For the candidate genes of the sex-combined meta-analysis, a univariate meta-regression analysis was performed using the meta package to examine the potential moderating effects of age, RNA integrity number (RIN), brain pH, and post-mortem interval (PMI) on the meta-analytical results.

## RESULTS

### 1. Included datasets

Overall, 14 independent GEO datasets were included. The process for selecting eligible datasets adhered to the PRISMA 2020 guidelines [57] (**Supplementary Fig. S1)**. We included 11 GEO datasets derived from brain tissues, totalizing 854 brain samples from 348 SZ individuals and 346 HC. Notably, some of the included GEO datasets provided data from several brain regions, thereby increasing the number of brain datasets from 11 to 14. Regarding peripheral blood samples, 3 GEO datasets were further included, totalizing 295 blood samples from 162 SZ individuals and 133 HC. The original studies from which the included datasets were obtained, the demographic and quality characteristics of the samples provided by the 17 included datasets are presented in **Supplementary Tables S1-S3**.

### 2. Differential expression meta-analysis in brain tissues

To analyze differences in gene expression irrespective of the brain region, a random-effect meta-analysis was performed on 13 to 14 studies, depending on the considered candidate gene, encompassing between 997 and 1,054 observations. In SZ individuals *vs*. HC, no significant differential expression was observed for *CSMD1* (SMD: 0.07 [−0.06; 0.20], adj-p=0.31), whereas a significant *CSMD2* overexpression was identified (SMD: 0.22 [0.05; 0.39], adj-p=0.026) (**Fig. 1**). Heterogeneity was negligible for *CSMD1* (I2=0.0%) and moderate for *CSMD2* (I2=40.8%). For these two genes, the Q-values indicated no significant variability between studies, the funnel plots and Egger’s tests indicated no presence of publication bias (**Supplementary Table S4**). Univariate meta-regression analyses revealed no statistically significant moderating effects of age, RIN, brain pH, or PMI on the effect sizes (**Supplementary Table S5**).

**Fig. 1.**
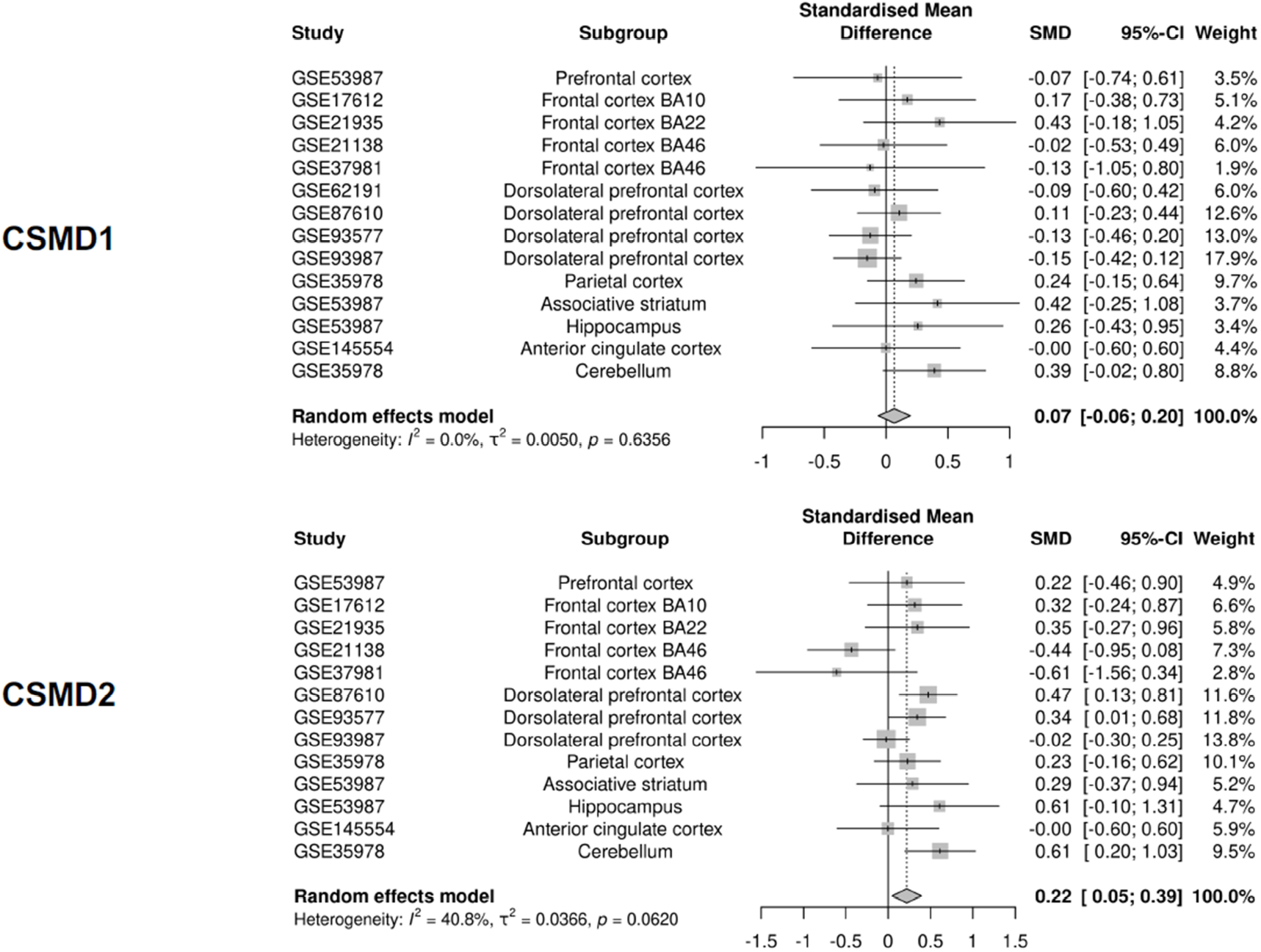
Sex-combined meta-analysis of differential expression of *CSMD1* and *CSMD2* genes in the brain tissues of individuals with schizophrenia vs. healthy controls

### 3. Sex-specific differential expression meta-analysis in brain tissues

To investigate the influence of sex on the meta-analytic results, separate analyses were conducted for female and male subgroups. To analyze differences in gene expression irrespective of the brain region, a random-effect meta-analysis was performed on 10 studies, depending on the considered candidate gene, encompassing between 189 and 321 observations. In the female subgroup, *CSMD2* overexpression reached nominal significance (SMD: 0.31 [0.02; 0.60], p=0.037) but did not remain significant after correction for multiple comparisons (adj-p=0.074). The random-effect meta-analysis revealed no significant change in the transcriptional level of *CSMD1* between SZ individuals and HC (SMD: 0.21 [−0.08; 0.50], adj-p=0.15) (**Fig. 2**). For these genes, the heterogeneity was low, the Q-values indicated no significant variability between studies, the funnel plots and Egger’s tests indicated no presence of publication bias (**Supplementary Table S6**). In the male subgroup, the random-effect meta-analysis revealed no significant change in the transcriptional level of *CSMD1* (SMD: 0.18 [−0.04; 0.40], adj-p=0.23) and *CSMD2* (SMD: 0.12 [−0.16; 0.41], adj-p=0.39) between SZ individuals and HC (**Fig. 3** and **Supplementary Table S7)**.

**Fig. 2.**
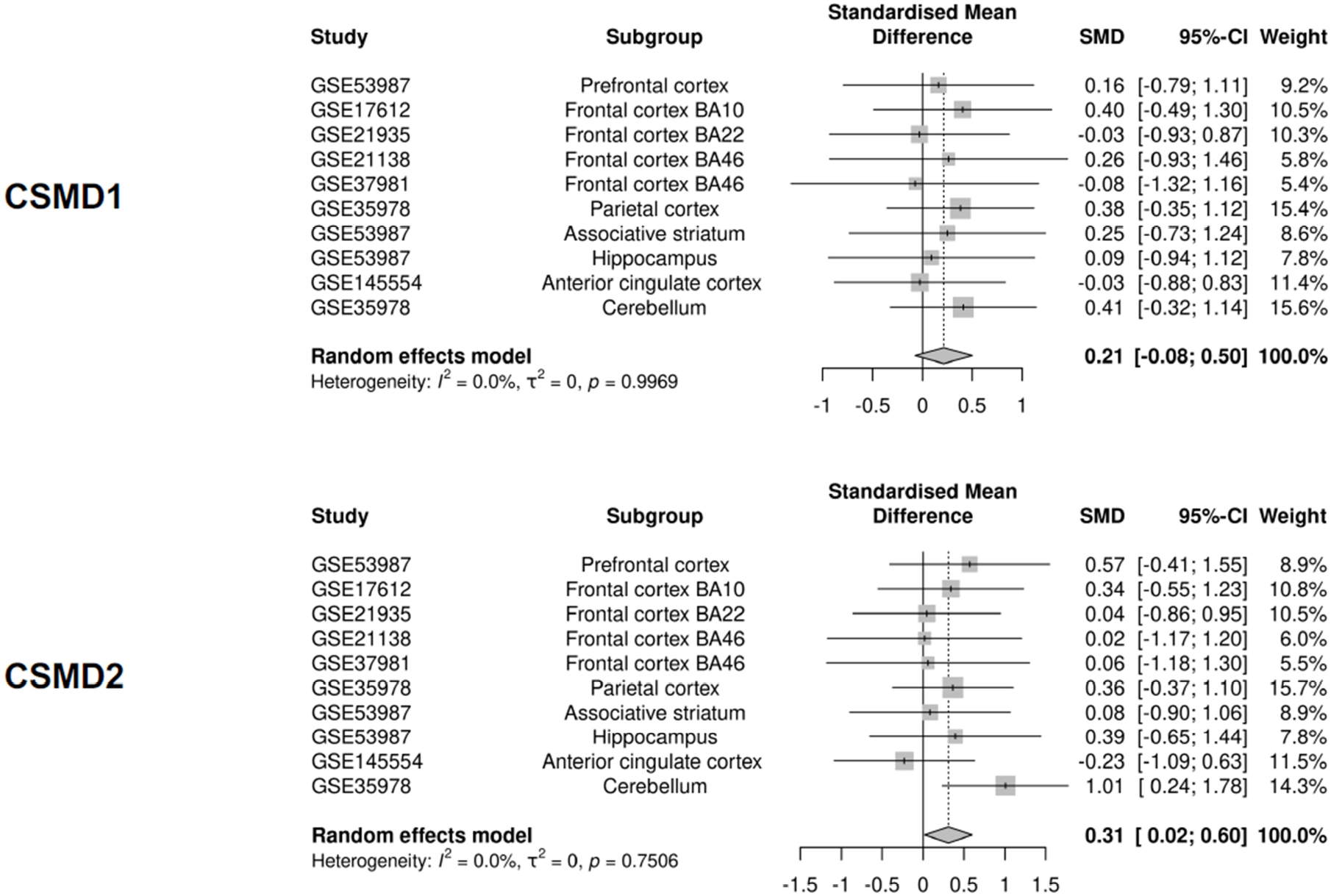
Meta-analysis of differential expression of *CSMD1* and *CSMD2* genes in the brain tissues of female subjects with schizophrenia vs. female healthy controls

**Fig. 3.**
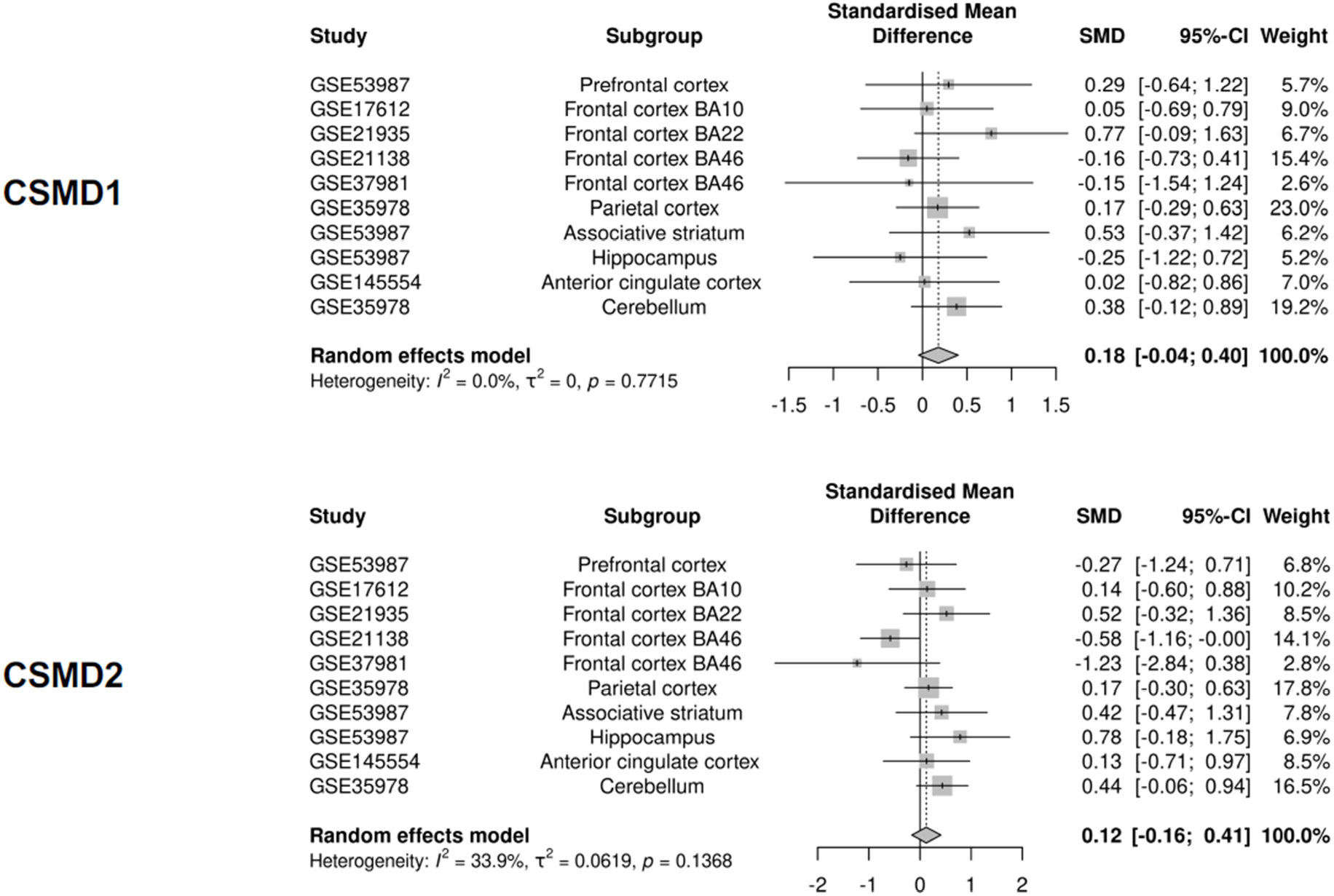
Meta-analysis of differential expression of *CSMD1* and *CSMD2* genes in the brain tissues of male subjects with schizophrenia vs. male healthy controls

### 4. Differential expression meta-analysis in peripheral blood

To analyze differences in gene expression, a random-effect meta-analysis was performed on 3 studies, encompassing 288 observations. The random-effect meta-analysis revealed no significant change in the transcriptional level of *CSMD1* (SMD: −0.19 [−0.59; 0.20], adj-p=0.52) and *CSMD2* (SMD: −0.08 [−0.31; 0.16], adj-p=0.52) between SZ individuals and HC (**Fig. 4** and **Supplementary Tables S8**).

**Fig. 4.**
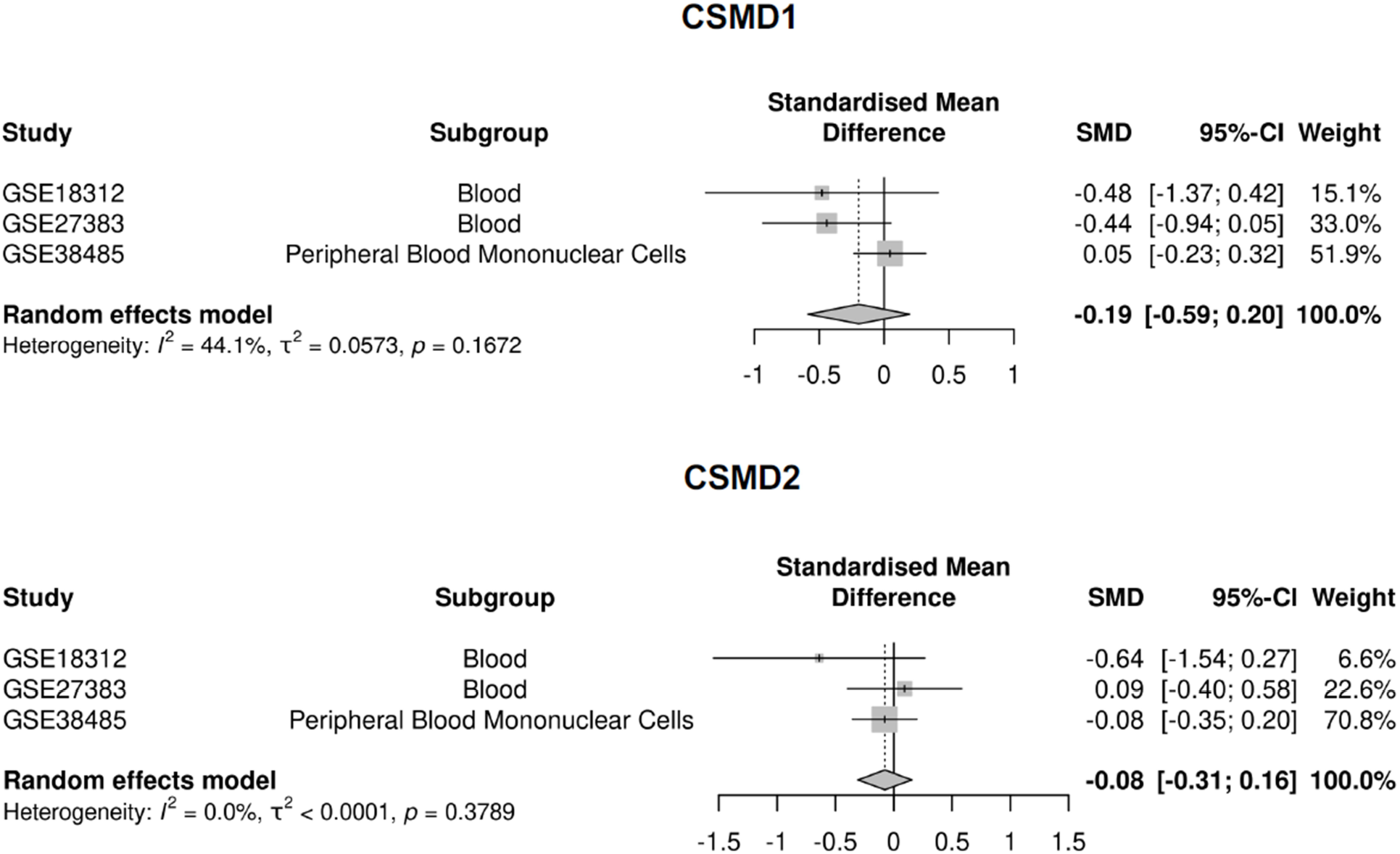
Sex-combined meta-analysis of differential expression of *CSMD1* and *CSMD2* genes in the peripheral blood of individuals with schizophrenia vs. healthy controls

## DISCUSSION

The present meta-analysis examined transcriptional alterations of the complement control protein-encoding genes *CSMD1* and *CSMD2* in SZ individuals compared with HC, using brain and blood-derived datasets, and including sex-stratified subgroup analyses. In brain tissues, schizophrenia was associated with a significant *CSMD2* overexpression, with a nominally significant increase also observed in the female-only meta-analysis. By contrast, no differential expression of either gene was detected in peripheral blood samples. These findings support the involvement of *CSMD2* in the neurobiological mechanisms underlying schizophrenia.

### Meta-analysis in the brain tissues

While GWAS [20] and subsequent candidate-gene studies conducted across populations of diverse ancestral backgrounds [27–29] have consistently reported an association between the rs10503253 variant in the complement control-related gene *CSMD1* and schizophrenia risk, no significant differential expression of *CSMD1* has been observed in SZ individuals compared with HC. At the brain level, these findings indicate that the genetic contribution of *CSMD1* to schizophrenia susceptibility may not involve alterations in its overall transcriptional levels.

This dissociation between genetic association with schizophrenia liability and transcriptional output may not extend to all members of the *CSMD* gene family, as evidenced by the *CSMD2* overexpression, which, to our knowledge, is reported here for the first time in the brain of SZ individuals compared with HC. In humans, *CSMD2* shows its highest expression in the brain and encodes a transmembrane protein that localizes to dendrites and synapses, where it is required for the development and maintenance of dendrites and dendritic spines [31]. Consistently, a voxelwise genome-wide association study identified *CSMD2* as a gene with high relevance to human brain structure [58], suggesting a role for *CSMD2* in synaptic development and function that may be perturbed in neuropsychiatric disorders [31]. In line with these results, common and/or rare variants in *CSMD2* have been associated with various neurodevelopmental disorders including ADHD [59, 60], autism [61], schizophrenia [29] and focal epilepsies [62]. *CSMD2* has further been associated with addiction vulnerability [63], alcohol dependence [64, 65] and comorbid depression [65], methamphetamine dependance and polysubstance abuse [66]. From a mechanistic perspective, the presence of CUB and Sushi domains, hallmark features of complement regulatory proteins, in all members of the *CSMD* gene family [67] has led to the hypothesis that CSMD2 protein contributes to the regulation of complement activation and inflammatory processes in the brain [30, 68]. Although the function of CSMD2 protein remains largely unexplored, it may share functional properties with its homolog CSMD1, which acts as an inhibitory regulator of complement-dependent synaptic elimination by protecting neurons from complement deposition and subsequent microglia-mediated engulfment of synaptic material [69, 70]. In the brain of SZ individuals, *CSMD2* overexpression may thus contribute to reduce the complement system activity, potentially reflecting a compensatory regulatory response to the well-documented overexpression of *C4* in schizophrenia [19, 21, 23, 24]. Further studies are required to determine whether this putative regulatory mechanism is sufficient to effectively counterbalance *C4* overexpression, or whether excessive complement system activity persists in the brain of SZ individuals. From another perspective, the temporal expression profile of *CSMD2* in the brain is characterized by a peak during neurodevelopment, supporting an essential role in brain maturation [62]. Accordingly, the persistence of *CSMD2* overexpression in the brain of adult individuals living with schizophrenia may represent either a failure in normal developmental transcriptional regulation or a potential reinduction in response to neuroinflammation, both of which could disrupt the fine-tuning of synaptic maintenance in schizophrenia. Overall, our results add to and refine the growing body of literature suggesting that schizophrenia pathogenesis may be partly driven by altered complement system activity [19, 26], by indicating that such aberrant complement activation may result from the combined effects of *C4* overexpression [21, 22, 24, 71] and altered *CSMD2* expression.

### Sexual dimorphism in the brain

In healthy adult human brains, large-scale transcriptomic analyses have revealed that numerous genes exhibit sex-dependent expression, including genes involved in immune-related functions [72]. Building on this baseline, our study examined pathological conditions and revealed nominally significant overexpression of *CSMD2* in brain tissue from females with schizophrenia as compared to healthy control females. These findings suggest that sexual dimorphism extends to the pathophysiology of schizophrenia and add to previous observations highlighting the complement system as a well-established contributor to sex differences in susceptibility to schizophrenia [73–75]. Sexual dimorphism in *CSMD2* brain expression may partly account for the observed differences in clinical presentation, disease progression, and severity between male and female individuals with schizophrenia [76].

### Meta-analysis in peripheral blood compartment

In the peripheral blood compartment, the present meta-analysis did not detect significant transcriptional differences between SZ individuals and HC. While no prior study has specifically examined *CSMD2* expression in schizophrenia, our findings did not replicate earlier reports regarding *CSMD1* transcription. Previous investigations in antipsychotic-naïve individuals experiencing a first episode of psychosis described reduced *CSMD1* expression, both in peripheral blood mononuclear cells (PBMCs) relative to HC [34] and in whole blood compared with healthy offspring of parents with schizophrenia [77]. Consistently, Liu et al. reported decreased *CSMD1* expression in PBMCs from antipsychotic-naïve individuals with schizophrenia vs. HC, followed by a significant increase after 12 weeks of antipsychotic treatment [33], suggesting that medication exposure may influence *CSMD1* transcription. The discrepancy between the present results and prior literature may reflect several methodological and clinical factors, including differences in biological material (PBMCs vs. whole blood, which are known to exhibit distinct expression profiles [78]), measurement techniques (RT-qPCR vs. microarray platforms), illness stage, treatment status and remission state (prior studies focused on untreated individuals with a short duration of illness). Importantly, although our meta-analysis included a substantially larger sample (n=288) than previous studies (n ranging from 80 to 160), potentially providing greater statistical power and a more stable estimate of peripheral transcriptional effects, the inclusion of more clinically heterogeneous subgroups may have obscured a *CSMD1* underexpression signal that could characterize a specific patient subgroup, in contrast to earlier smaller studies that examined more homogeneous populations.

In this study, transcriptional alterations observed in peripheral blood did not parallel those detected in brain tissue from SZ individuals. The lack of concordant brain-blood changes for *CSMD1* and *CSMD2* supports the view that peripheral gene expression may not reliably reflect brain transcriptional processes, consistent with prior concerns [36]. This discrepancy may also be influenced by methodological and biological factors, including the fact that brain and peripheral samples were obtained from different individuals, as well as the possibility that schizophrenia-related alterations may differentially affect brain and peripheral tissues [36, 79].

### Limits

The interpretation of the present findings should be considered in light of several limitations. First, the included datasets were heterogeneous in terms of brain regions and patient characteristics, which may obscure region- or subgroup-specific effects. The number of available blood datasets was limited, reducing statistical power. Sex-stratified analyses were restricted to subsets of studies with annotated demographic data, potentially underestimating sex-specific patterns. Second, multiple clinical and biological factors - including antipsychotic exposure, cause of death, history of suicide attempts, somatic comorbidities, illness chronicity, and symptom profiles - may have confounded the results. Owing to incomplete reporting across the included studies, it was not possible to fully adjust for all potential confounders in supplementary analyses. Third, several datasets reported significant differences between SZ individuals and HC in RNA integrity number (RIN), brain pH, and postmortem interval (PMI), variables that could influence transcriptional measurements. However, meta-regression analyses did not identify these factors as significant moderators, supporting the robustness of the findings (**Supplementary Table S5**). Fourth, all datasets were generated using microarray technology, and the present results are therefore subject to inherent methodological constraints. These include limitations in detecting low-abundance transcripts due to background hybridization, potential probe-related specificity issues arising from cross- or non-specific hybridization, and the restriction to genes represented on the array platform. Fifth, postmortem studies provide only a static snapshot of neurobiology at the time of death and cannot capture the dynamic alterations occurring at disease onset. This is a key limitation in schizophrenia, whose pathophysiology is rooted in early neurodevelopment. This consideration is particularly relevant given that *CSMD1* and *CSMD2* exhibit developmentally regulated brain expression, with both genes predominantly expressed during early stages, and *CSMD2* showing especially high expression in the developing brain [62]. In contrast, the brains analyzed in the present study were obtained from adults aged 42 to 73 years. In the brain of SZ individuals, our findings therefore suggest that complement system alterations are observed well beyond adolescence and the period of synaptic pruning. However, the cross-sectional nature of postmortem data prevents us from determining whether the observed *CSMD2* transcriptional changes originate before or during adolescence (possibly impacting the synaptic pruning process), or arise later, potentially as a compensatory response to the well-documented cerebral inflammation associated with schizophrenia.

## Conclusion

The present findings highlight a dissociation between genetic vulnerability and transcriptional output within the *CSMD* gene family in schizophrenia. Although *CSMD1* is genetically associated with disease risk, this liability does not appear to translate into detectable changes in its overall brain expression, whereas *CSMD2* shows a selective overexpression, reported here for the first time in the brain tissues of individuals with schizophrenia. The absence of parallel alterations in peripheral blood further indicates that peripheral transcriptional measures may not reliably capture central molecular processes. Taken together, these results strengthen the view of the complement system as a convergent biological pathway in schizophrenia and suggest that aberrant complement activity may arise from the combined influence of *C4* and *CSMD2* overexpression. This integrated perspective contributes to a more nuanced framework linking immune-related molecular dysregulation to schizophrenia pathophysiology. Future research should clarify the spatio-temporal dynamics and functional consequences of these transcriptional changes, particularly during neurodevelopmental windows critical for synaptic refinement, and determine how complement-related dysregulation contributes to disease mechanisms.

## REFERENCES

1. Tandon R, Keshavan MS, Nasrallah HA. Schizophrenia, ‘just the facts’ what we know in 2008. 2. Epidemiology and etiology. Schizophr Res. 2008;102:1–18.

2. Owen MJ, Sawa A, Mortensen PB. Schizophrenia. Lancet. 2016;388:86–97.

3. Laursen TM, Nordentoft M, Mortensen PB. Excess early mortality in schizophrenia. Annu Rev Clin Psychol. 2014;10:425–448.

4. Al-Diwani AAJ, Pollak TA, Irani SR, Lennox BR. Psychosis: an autoimmune disease? Immunology. 2017;152:388–401.

5. Sullivan PF, Kendler KS, Neale MC. Schizophrenia as a complex trait: evidence from a meta-analysis of twin studies. Arch Gen Psychiatry. 2003;60:1187–1192.

6. Birnbaum R, Weinberger DR. Genetic insights into the neurodevelopmental origins of schizophrenia. Nat Rev Neurosci. 2017;18:727–740.

7. Johnson MB, Stevens B. Pruning hypothesis comes of age. Nature. 2018;554:438–439.

8. Rocco BR, DeDionisio AM, Lewis DA, Fish KN. Alterations in a Unique Class of Cortical Chandelier Cell Axon Cartridges in Schizophrenia. Biol Psychiatry. 2017;82:40–48.

9. Berdenis van Berlekom A, Muflihah CH, Snijders GJLJ, MacGillavry HD, Middeldorp J, Hol EM, et al. Synapse Pathology in Schizophrenia: A Meta-analysis of Postsynaptic Elements in Postmortem Brain Studies. Schizophrenia Bulletin. 2020;46:374–386.

10. Mallya AP, Deutch AY. (Micro)Glia as Effectors of Cortical Volume Loss in Schizophrenia. Schizophrenia Bulletin. 2018;44:948–957.

11. Selemon LD, Goldman-Rakic PS.The reduced neuropil hypothesis: a circuit based model of schizophrenia. Biol Psychiatry. 1999;45:17–25.

12. Schultz CC, Koch K, Wagner G, Roebel M, Nenadic I, Schachtzabel C, et al. Complex pattern of cortical thinning in schizophrenia: Results from an automated surface based analysis of cortical thickness. Psychiatry Research: Neuroimaging. 2010;182:134–140.

13. Jung WH, Kim JS, Jang JH, Choi J-S, Jung MH, Park J-Y, et al. Cortical thickness reduction in individuals at ultra-high-risk for psychosis. Schizophr Bull. 2011;37:839–849.

14. Schultz CC, Koch K, Wagner G, Roebel M, Schachtzabel C, Gaser C, et al. Reduced cortical thickness in first episode schizophrenia. Schizophrenia Research. 2010;116:204–209.

15. Magdalon J, Mansur F, Teles E Silva AL, de Goes VA, Reiner O, Sertié AL. Complement System in Brain Architecture and Neurodevelopmental Disorders. Front Neurosci. 2020;14:23.

16. Presumey J, Bialas AR, Carroll MC. Complement System in Neural Synapse Elimination in Development and Disease. Adv Immunol. 2017;135:53–79.

17. Schafer DP, Lehrman EK, Kautzman AG, Koyama R, Mardinly AR, Yamasaki R, et al. Microglia sculpt postnatal neural circuits in an activity and complement-dependent manner. Neuron. 2012;74:691–705.

18. Stevens B, Allen NJ, Vazquez LE, Howell GR, Christopherson KS, Nouri N, et al. The classical complement cascade mediates CNS synapse elimination. Cell. 2007;131:1164– 1178.

19. Schizophrenia Working Group of the Psychiatric Genomics Consortium, Sekar A, Bialas AR, de Rivera H, Davis A, Hammond TR, et al. Schizophrenia risk from complex variation of complement component 4. Nature. 2016;530:177–183.

20. Schizophrenia Working Group of the Psychiatric Genomics Consortium. Biological insights from 108 schizophrenia-associated genetic loci. Nature. 2014;511:421–427.

21. Gandal MJ, Zhang P, Hadjimichael E, Walker RL, Chen C, Liu S, et al. Transcriptome-wide isoform-level dysregulation in ASD, schizophrenia, and bipolar disorder. Science. 2018;362.

22. Fromer M, Roussos P, Sieberts SK, Johnson JS, Kavanagh DH, Perumal TM, et al. Gene expression elucidates functional impact of polygenic risk for schizophrenia. Nat Neurosci. 2016;19:1442–1453.

23. Gandal MJ, Haney JR, Parikshak NN, Leppa V, Ramaswami G, Hartl C, et al. Shared molecular neuropathology across major psychiatric disorders parallels polygenic overlap. Science. 2018;359:693–697.

24. Rey R, Suaud-Chagny M-F, Bohec A-L, Dorey J-M, d’Amato T, Tamouza R, et al. Overexpression of complement component c4 in the dorsolateral prefrontal cortex, parietal cortex, superior temporal gyrus and associative striatum of patients with schizophrenia. Brain Behav Immun. 2020. 19 August 2020. 10.1016/j.bbi.2020.08.019.

25. Prasad KM, Chowdari KV, D’Aiuto LA, Iyengar S, Stanley JA, Nimgaonkar VL. Neuropil contraction in relation to Complement C4 gene copy numbers in independent cohorts of adolescent-onset and young adult-onset schizophrenia patients-a pilot study. Transl Psychiatry. 2018;8:134.

26. Woo JJ, Pouget JG, Zai CC, Kennedy JL. The complement system in schizophrenia: where are we now and what’s next? Mol Psychiatry. 2020;25:114–130.

27. Mihoub O, Ben Chaaben A, Boukouaci W, Lajnef M, Ayari F, El Kefi H, et al. CSMD1 rs10503253 increases schizophrenia risk in a Tunisian population-group. Encephale. 2024;50:380–385.

28. Liu W, Liu F, Xu X, Bai Y. Replicated association between the European GWAS locus rs10503253 at CSMD1 and schizophrenia in Asian population. Neurosci Lett. 2017;647:122–128.

29. Håvik B, Le Hellard S, Rietschel M, Lybæk H, Djurovic S, Mattheisen M, et al. The complement control-related genes CSMD1 and CSMD2 associate to schizophrenia. Biol Psychiatry. 2011;70:35–42.

30. Kraus DM, Elliott GS, Chute H, Horan T, Pfenninger KH, Sanford SD, et al. CSMD1 is a novel multiple domain complement-regulatory protein highly expressed in the central nervous system and epithelial tissues. J Immunol. 2006;176:4419–4430.

31. Gutierrez MA, Dwyer BE, Franco SJ. Csmd2 Is a Synaptic Transmembrane Protein that Interacts with PSD-95 and Is Required for Neuronal Maturation. eNeuro. 2019;6.

32. Baum ML, Wilton DK, Fox RG, Carey A, Hsu Y-HH, Hu R, et al. CSMD1 regulates brain complement activity and circuit development. Brain Behav Immun. 2024;119:317–332.

33. Liu Y, Fu X, Tang Z, Li C, Xu Y, Zhang F, et al. Altered expression of the CSMD1 gene in the peripheral blood of schizophrenia patients. BMC Psychiatry. 2019;19:113.

34. Hatzimanolis A, Foteli S, Stefanatou P, Ntigrintaki A-A, Ralli I, Kollias K, et al. Deregulation of complement components C4A and CSMD1 peripheral expression in first-episode psychosis and links to cognitive ability. Eur Arch Psychiatry Clin Neurosci. 2022;272:1219–1228.

35. Abd El Gayed EM, Rizk MS, Ramadan AN, Bayomy NR. mRNA Expression of the CUB and Sushi Multiple Domains 1 (CSMD1) and Its Serum Protein Level as Predictors for Psychosis in the Familial High-Risk Children and Young Adults. ACS Omega. 2021;6:24128–24138.

36. Cai C, Langfelder P, Fuller TF, Oldham MC, Luo R, van den Berg LH, et al. Is human blood a good surrogate for brain tissue in transcriptional studies? BMC Genomics. 2010;11:589.

37. Wu X, Shukla R, Alganem K, Zhang X, Eby HM, Devine EA, et al. Transcriptional profile of pyramidal neurons in chronic schizophrenia reveals lamina-specific dysfunction of neuronal immunity. Mol Psychiatry. 2021;26:7699–7708.

38. Maycox PR, Kelly F, Taylor A, Bates S, Reid J, Logendra R, et al. Analysis of gene expression in two large schizophrenia cohorts identifies multiple changes associated with nerve terminal function. Mol Psychiatry. 2009;14:1083–1094.

39. Narayan S, Tang B, Head SR, Gilmartin TJ, Sutcliffe JG, Dean B, et al. Molecular profiles of schizophrenia in the CNS at different stages of illness. Brain Res. 2008;1239:235–248.

40. Barnes MR, Huxley-Jones J, Maycox PR, Lennon M, Thornber A, Kelly F, et al. Transcription and pathway analysis of the superior temporal cortex and anterior prefrontal cortex in schizophrenia. J Neurosci Res. 2011;89:1218–1227.

41. Chen C, Cheng L, Grennan K, Pibiri F, Zhang C, Badner JA, et al. Two gene co-expression modules differentiate psychotics and controls. Mol Psychiatry. 2013;18:1308–1314.

42. Pietersen CY, Mauney SA, Kim SS, Lim MP, Rooney RJ, Goldstein JM, et al. Molecular profiles of pyramidal neurons in the superior temporal cortex in schizophrenia. J Neurogenet. 2014;28:53–69.

43. Lanz TA, Reinhart V, Sheehan MJ, Rizzo SJS, Bove SE, James LC, et al. Postmortem transcriptional profiling reveals widespread increase in inflammation in schizophrenia: a comparison of prefrontal cortex, striatum, and hippocampus among matched tetrads of controls with subjects diagnosed with schizophrenia, bipolar or major depressive disorder. Transl Psychiatry. 2019;9:151.

44. de Baumont A, Maschietto M, Lima L, Carraro DM, Olivieri EH, Fiorini A, et al. Innate immune response is differentially dysregulated between bipolar disease and schizophrenia. Schizophr Res. 2015;161:215–221.

45. Arion D, Huo Z, Enwright JF, Corradi JP, Tseng G, Lewis DA. Transcriptome Alterations in Prefrontal Pyramidal Cells Distinguish Schizophrenia From Bipolar and Major Depressive Disorders. Biol Psychiatry. 2017;82:594–600.

46. Datta D, Enwright JF, Arion D, Paspalas CD, Morozov YM, Lewis DA, et al. Mapping Phosphodiesterase 4D (PDE4D) in Macaque Dorsolateral Prefrontal Cortex: Postsynaptic Compartmentalization in Layer III Pyramidal Cell Circuits. Front Neuroanat. 2020;14:578483.

47. Arion D, Corradi JP, Tang S, Datta D, Boothe F, He A, et al. Distinctive transcriptome alterations of prefrontal pyramidal neurons in schizophrenia and schizoaffective disorder. Mol Psychiatry. 2015;20:1397–1405.

48. Bousman CA, Chana G, Glatt SJ, Chandler SD, Lucero GR, Tatro E, et al. Preliminary evidence of ubiquitin proteasome system dysregulation in schizophrenia and bipolar disorder: convergent pathway analysis findings from two independent samples. Am J Med Genet B Neuropsychiatr Genet. 2010;153B:494–502.

49. van Beveren NJM, Buitendijk GHS, Swagemakers S, Krab LC, Röder C, de Haan L, et al. Marked reduction of AKT1 expression and deregulation of AKT1-associated pathways in peripheral blood mononuclear cells of schizophrenia patients. PLoS ONE. 2012;7:e32618.

50. de Jong S, Boks MPM, Fuller TF, Strengman E, Janson E, de Kovel CGF, et al. A gene co-expression network in whole blood of schizophrenia patients is independent of antipsychotic-use and enriched for brain-expressed genes. PLoS ONE. 2012;7:e39498.

51. Dumas J, Gargano MA, Dancik GM. shinyGEO: a web-based application for analyzing gene expression omnibus datasets. Bioinformatics. 2016;32:3679–3681.

52. Davis S, Meltzer PS. GEOquery: a bridge between the Gene Expression Omnibus (GEO) and BioConductor. Bioinformatics. 2007;23:1846–1847.

53. Schwarzer G, Carpenter JR, Rücker G. Meta-Analysis with R. Cham: Springer International Publishing; 2015.

54. Higgins JPT, Thompson SG, Spiegelhalter DJ. A re-evaluation of random-effects meta-analysis. J R Stat Soc Ser A Stat Soc. 2009;172:137–159.

55. Higgins JPT, Thompson SG, Deeks JJ, Altman DG. Measuring inconsistency in meta-analyses. BMJ. 2003;327:557–560.

56. Egger M, Smith GD, Schneider M, Minder C. Bias in meta-analysis detected by a simple, graphical test. BMJ. 1997;315:629–634.

57. Page MJ, McKenzie JE, Bossuyt PM, Boutron I, Hoffmann TC, Mulrow CD, et al. The PRISMA 2020 statement: an updated guideline for reporting systematic reviews. BMJ. 2021:71.

58. Stein JL, Hua X, Lee S, Ho AJ, Leow AD, Toga AW, et al. Voxelwise genome-wide association study (vGWAS). Neuroimage. 2010;53:1160–1174.

59. Lesch K-P, Timmesfeld N, Renner TJ, Halperin R, Röser C, Nguyen TT, et al. Molecular genetics of adult ADHD: converging evidence from genome-wide association and extended pedigree linkage studies. J Neural Transm (Vienna). 2008;115:1573–1585.

60. Kim DS, Burt AA, Ranchalis JE, Wilmot B, Smith JD, Patterson KE, et al. Sequencing of sporadic Attention-Deficit Hyperactivity Disorder (ADHD) identifies novel and potentially pathogenic de novo variants and excludes overlap with genes associated with autism spectrum disorder. Am J Med Genet B Neuropsychiatr Genet. 2017;174:381–389.

61. Khan H, Harripaul R, Mikhailov A, Herzi S, Bowers S, Ayub M, et al. Biallelic variants identified in 36 Pakistani families and trios with autism spectrum disorder. Sci Rep. 2024;14:9230.

62. Li X, Wang L, Liang X-Y, Zhang H-W, Shi J-G, Guo J, et al. Variants in CSMD2 and CSMD3, genes involved in synaptogenesis, are associated with epilepsies. Epilepsia. 2025;66:4394–4410.

63. Liu Q-R, Drgon T, Johnson C, Walther D, Hess J, Uhl GR. Addiction molecular genetics: 639,401 SNP whole genome association identifies many ‘cell adhesion’ genes. Am J Med Genet B Neuropsychiatr Genet. 2006;141B:918–925.

64. Johnson C, Drgon T, Liu Q-R, Walther D, Edenberg H, Rice J, et al. Pooled association genome scanning for alcohol dependence using 104,268 SNPs: validation and use to identify alcoholism vulnerability loci in unrelated individuals from the collaborative study on the genetics of alcoholism. Am J Med Genet B Neuropsychiatr Genet. 2006;141B:844– 853.

65. Edwards AC, Aliev F, Bierut LJ, Bucholz KK, Edenberg H, Hesselbrock V, et al. Genome-wide association study of comorbid depressive syndrome and alcohol dependence. Psychiatr Genet. 2012;22:31–41.

66. Uhl GR, Drgon T, Johnson C, Fatusin OO, Liu Q-R, Contoreggi C, et al. ‘Higher order’ addiction molecular genetics: convergent data from genome-wide association in humans and mice. Biochem Pharmacol. 2008;75:98–111.

67. Lau WL, Scholnick SB. Identification of two new members of the CSMD gene family. Genomics. 2003;82:412–415.

68. Athanasiu L, Giddaluru S, Fernandes C, Christoforou A, Reinvang I, Lundervold AJ, et al. A genetic association study of CSMD1 and CSMD2 with cognitive function. Brain Behav Immun. 2017;61:209–216.

69. Byrne RAJ, Nimmo J, Torvell M, Carpanini SM, Daskoulidou N, Hughes TR, et al. The schizophrenia-associated gene CSMD1 encodes a complement classical pathway inhibitor predominantly expressed by astrocytes and at synapses in mice and humans. Brain Behav Immun. 2025;127:287–302.

70. Baum ML, Wilton DK, Fox RG, Carey A, Hsu Y-HH, Hu R, et al. CSMD1 regulates brain complement activity and circuit development. Brain Behav Immun. 2024;119:317–332.

71. Pérez-Santiago J, Diez-Alarcia R, Callado LF, Zhang JX, Chana G, White CH, et al. A combined analysis of microarray gene expression studies of the human prefrontal cortex identifies genes implicated in schizophrenia. J Psychiatr Res. 2012;46:1464–1474.

72. He S, Zhang X, Zhu H. Human-specific protein-coding and lncRNA genes cast sex-biased genes in the brain and their relationships with brain diseases. Biol Sex Differ. 2024;15:86.

73. Kamitaki N, Sekar A, Handsaker RE, de Rivera H, Tooley K, Morris DL, et al. Complement genes contribute sex-biased vulnerability in diverse disorders. Nature. 2020;582:577– 581.

74. Fournier LA, Phadke RA, Salgado M, Brack A, Nocon JC, Bolshakova S, et al. Overexpression of the schizophrenia risk gene C4 in PV cells drives sex-dependent behavioral deficits and circuit dysfunction. iScience. 2024;27:110800.

75. Hui CW, Vecchiarelli HA, Gervais É, Luo X, Michaud F, Scheefhals L, et al. Sex Differences of Microglia and Synapses in the Hippocampal Dentate Gyrus of Adult Mouse Offspring Exposed to Maternal Immune Activation. Front Cell Neurosci. 2020;14:558181.

76. Teymouri K, Ebrahimi M, Chen CC, Sriretnakumar V, Mohiuddin AG, Tiwari AK, et al. Sex-dependent association study of complement C4 gene with treatment-resistant schizophrenia and hospitalization frequency. Psychiatry Res. 2024;342:116202.

77. Abd El Gayed EM, Rizk MS, Ramadan AN, Bayomy NR. mRNA Expression of the CUB and Sushi Multiple Domains 1 (CSMD1) and Its Serum Protein Level as Predictors for Psychosis in the Familial High-Risk Children and Young Adults. ACS Omega. 2021;6:24128–24138.

78. Gautam A, Donohue D, Hoke A, Miller SA, Srinivasan S, Sowe B, et al. Investigating gene expression profiles of whole blood and peripheral blood mononuclear cells using multiple collection and processing methods. PLoS ONE. 2019;14:e0225137.

79. Rey R, Suaud-Chagny M-F, Dorey J-M, Teyssier J-R, d’Amato T. Widespread transcriptional disruption of the microRNA biogenesis machinery in brain and peripheral tissues of individuals with schizophrenia. Transl Psychiatry. 2020;10:376.

