## Supplementary materials for "Dissociation Between Genetic Risk and Transcriptional Output in Schizophrenia: A Cross-Tissue Meta-Analysis of *CSMD1* and *CSMD2* Expression"

Address: Le Vinatier - Psychiatrie Universitaire Lyon Métropole, Pôle EST - Centre expert schizophrénie de Lyon, 95 boulevard Pinel BP 30039, 69678 Bron Cedex, France.

ORCID ID: 0000-0002-4603-3575

**Supplementary Fig. S1.** Flow Diagram of included datasets

**Supplementary Table S1.** Original studies from which the included datasets were obtained

**Supplementary Table S2.** Demographic and quality characteristics of the postmortem brain samples

**Supplementary Table S3.** Demographic and quality characteristics of the peripheral blood samples

**Supplementary Table S4.** Results of the differential expression meta-analysis in brain tissues

**Supplementary Table S5.** QM statistics from univariate meta-regression analysis of moderators (age, pH, PMI, RIN) performed for genes differentially expressed in the sex-combined meta-analysis

**Supplementary Table S6.** Results of the differential expression meta-analysis in the brain tissues of female subjects

**Supplementary Table S7.** Results of the differential expression meta-analysis in the brain tissues of male subjects

**Supplementary Table S8.** Results of the differential expression meta-analysis in peripheral blood

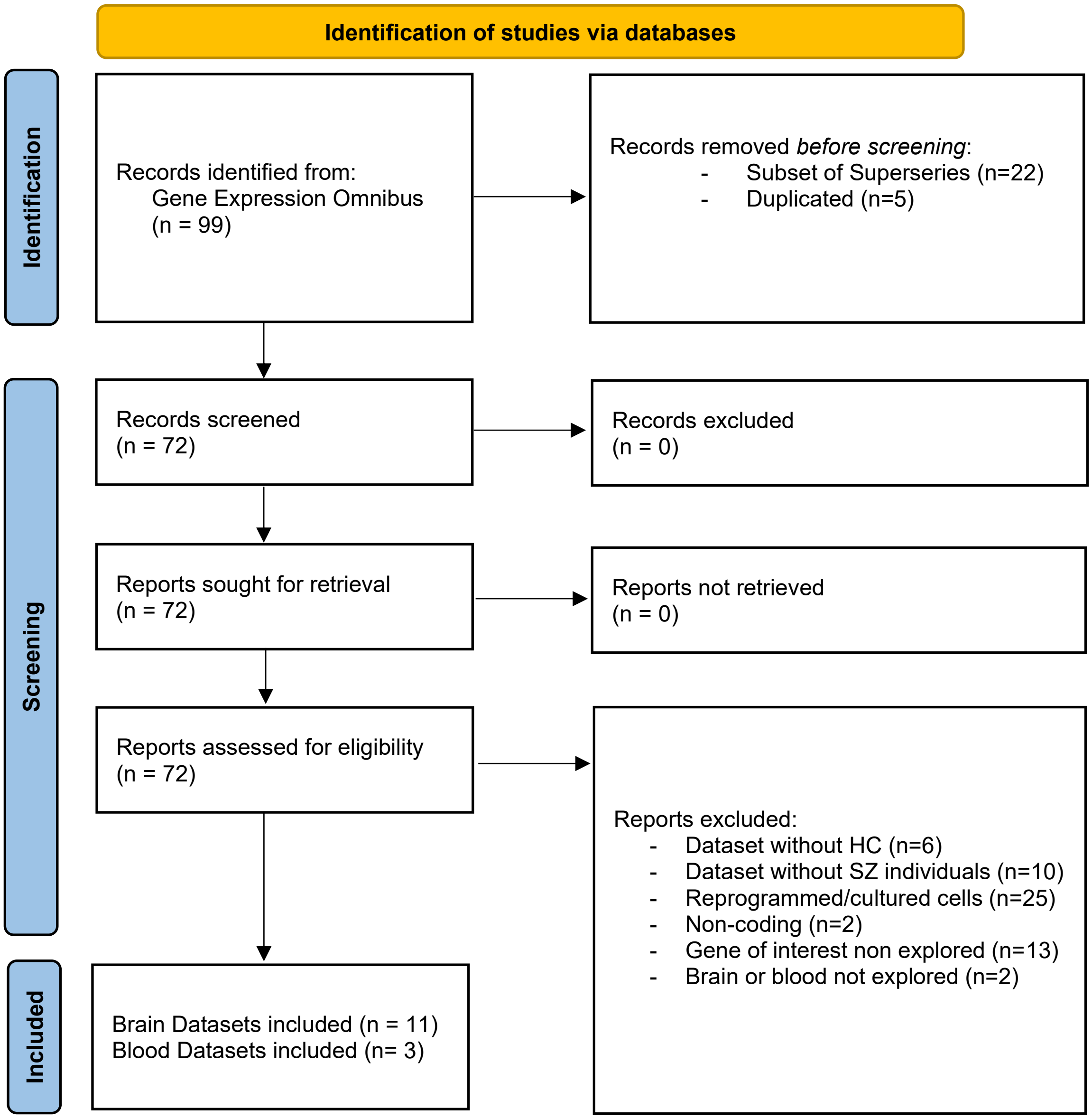


**Supplementary Fig. S1.** Flow Diagram of included datasets

**Supplementary Table S1.** Original studies from which the included datasets were obtained

| GEO accession | Tissue | Region / Cell type | Microarray platform | Year | n subjects healthy controls Total/F/M | n subjects SZ individuals Total/F/M | PubMed ID |
|---|---|---|---|---|---|---|---|
| **BRAIN DATASETS** | | | | | | | |
| **GSE145554** | Brain | Anterior cingulate cortex (superficial and deep pyramidal neurons) | [HuGene-1_0-st] Affymetrix Human Gene 1.0 ST Array [transcript (gene) version] | 2020 | 23/11/12 | 20/10/10 | 34272489 |
| **GSE17612** | Brain | Prefrontal cortex (BA10) | GPL570 [HG-U133_Plus_2] Affymetrix Human Genome U133 Plus 2.0 Array | 2009 | 23/11/11 (sex data missing for one sample) | 28/9/19 | 19255580 |
| **GSE21138** | Brain | Prefrontal cortex (BA46) | GPL570 [HG-U133_Plus_2] Affymetrix Human Genome U133 Plus 2.0 Array | 2010 | 29/5/24 | 30/6/24 | 18778695 |
| **GSE21935** | Brain | Superior temporal cortex (BA22) | GPL570 [HG-U133_Plus_2] Affymetrix Human Genome U133 Plus 2.0 Array | 2011 | 19/9/10 | 23/10/13 | 21538462 |
| **GSE35978** | Brain | Cerebellum | GPL6244 [HuGene-1_0-st] Affymetrix Human Gene 1.0 ST Array [transcript (gene) version] | 2012 | 44/12/32 | 50/19/31 | 23147385 |
| | Brain | Parietal cortex | GPL6244 [HuGene-1_0-st] Affymetrix Human Gene 1.0 ST Array [transcript (gene) version] | 2012 | 50/15/35 | 51/14/37 | |
| **GSE37981** | Brain | Superior temporal gyrus (pyramidal cells in layer III) | GPL1352 [U133_X3P] Affymetrix Human X3P Array | 2012 | 9/5/4 | 9/5/4 | 24702465 |
| **GSE53987** | Brain | Prefrontal cortex (BA46) | GPL570 [HG-U133_Plus_2] Affymetrix Human Genome U133 Plus 2.0 Array | 2014 | 19/9/10 | 15/8/7 | 31123247 |
| | Brain | Associative striatum | GPL570 [HG-U133_Plus_2] Affymetrix Human Genome U133 Plus 2.0 Array | 2014 | 18/8/10 | 18/8/10 | |
| | Brain | Hippocampus | GPL570 [HG-U133_Plus_2] Affymetrix Human Genome U133 Plus 2.0 Array | 2014 | 18/9/9 | 15/6/9 | |
| **GSE62191** | Brain | Frontal Cortex | GPL4133 Agilent-014850 Whole Human Genome Microarray 4x44K G4112F | 2014 | 30/7/23 | 29/6/23 | 25487697 |
| **GSE87610** | Brain | Dorsolateral prefrontal cortex (DLPFC) | GPL13667 [HG-U219] Affymetrix Human Genome U219 Array | 2017 | 19/9/10 | 19/9/10 | 28476208 |
| **GSE93987** | Brain | Dorsolateral prefrontal cortex (DLPFC) | GPL13158 [HT_HG-U133_Plus_PM] Affymetrix HT HG-U133+ PM Array Plate | 2014 | 106/NC/NC | 102/NC/NC | 25560755 |
| **GSE93577** | Brain | Dorsolateral prefrontal cortex (DLPFC) | GPL13667 [HG-U219] Affymetrix Human Genome U219 Array | 2018 | 19/9/10 | 19/9/10 | 33328902 |

| PERIPHERAL BLOOD DATASETS | | | | | | | |
|---|---|---|---|---|---|---|---|
| **GSE18312** | Blood | Blood | gpl5175 [HuEx-1_0-st] Affymetrix Human Exon 1.0 ST Array [transcript (gene) version] | 2009 | 8/3/5 | 13/4/9 | 19582768 |
| **GSE27383** | Blood | Peripheral Blood Mononuclear Cells (PBMC) | GPL570 [HG-U133_Plus_2] Affymetrix Human Genome U133 Plus 2.0 Array | 2013 | 29/0/29 | 43/0/43 | 22393424 |
| **GSE38485** | Blood | Whole blood | GPL6883 Illumina HumanRef-8 v3.0 expression beadchip | 2012 | 96/54/42 | 106/30/76 | 22761806 |

**Supplementary Table S2.** Demographic and quality characteristics of the postmortem brain samples

| Study | Brain Region | n subjects Healthy controls | n subjects SZ individuals | Age (y, mean ± SD) | | | Gender (F/M) | | | RIN (mean ± SD) | | | pH (mean ± SD) | | | PMI (hrs, mean ± SD) | | |
|---|---|---|---|---|---|---|---|---|---|---|---|---|---|---|---|---|---|---|
| | | | | Healthy controls | SZ individuals | p | Healthy controls | SZ individuals | p | Healthy controls | SZ individuals | p | Healthy controls | SZ individuals | p | Healthy controls | SZ individuals | p |
| **GSE145554** | Anterior cingulate cortex | 23 | 20 | 73,52±10,76 | 73,05±9,98 | 0,88 | 11/12 | 10/10 | 1 | NC/NC | NC/NC | - | 6,43±0,26 | 6,32±0,43 | 0,33 | 6,28±3,66 | 6,8±3,09 | 0,63 |
| **GSE17612** | Prefrontal cortex (BA10) | 23 | 28 | 69,04±21,55 | 73,32±15,2 | 0,43 | 11/11 | 9/19 | 0,25 | NC/NC | NC/NC | - | 6,5±0,29 | 6,15±0,21 | **$2,1.10^{-5}$** | 9,9±4,39 | 8,71±6,98 | 0,49 |
| **GSE21138** | Prefrontal cortex (BA46) | 29 | 30 | 44,72±16,14 | 43,4±16,96 | 0,76 | 5/24 | 6/24 | 0,42 | NC/NC | NC/NC | - | 6,31±0,17 | 6,24±0,22 | 0,18 | NC/NC | NC/NC | - |
| **GSE21935** | Superior temporal cortex (BA22) | 19 | 23 | 67,68±22,24 | 72,17±16,93 | 0,47 | 9/10 | 10/13 | 1 | NC/NC | NC/NC | - | 6,49±0,32 | 6,16±0,17 | **$4,1.10^{-4}$** | 9,11±4,33 | 7,13±5,75 | 0,22 |
| **GSE35978** | Cerebellum | 44 | 50 | 45,8±9,33 | 43,18±9,53 | 0,18 | 12/32 | 19/31 | 0,83 | NC/NC | NC/NC | - | 6,47±0,32 | 6,42±0,25 | 0,50 | NC/NC | NC/NC | - |
| | Parietal Cortex | 50 | 51 | 45,5±8,99 | 42,64±9,87 | 0,13 | 15/35 | 14/37 | 0,28 | NC/NC | NC/NC | - | 6,51±0,31 | 6,37±0,29 | **0,021** | NC/NC | NC/NC | - |
| **GSE37981** | Superior temporal gyrus | 9 | 9 | 69,11±20,57 | 67,11±19,8 | 0,83 | 5/4 | 5/4 | 1 | NC/NC | NC/NC | - | 6,46±0,12 | 6,34±0,11 | **0,042** | 16,71±1,6 | 16,9±1,85 | 0,82 |
| **GSE53987** | Prefrontal cortex (BA46) | 19 | 15 | 48,05±10,64 | 46±8,64 | 0,54 | 9/10 | 8/7 | 1 | 7,85±0,62 | 7,64±0,68 | 0,36 | 6,59±0,21 | 6,53±0,39 | 0,60 | 19,53±5,09 | 18,91±6,69 | 0,77 |
| | Associative striatum | 18 | 18 | 48,44±10,82 | 45±8,75 | 0,31 | 8/10 | 8/10 | 1 | 8,21±0,69 | 7,87±0,79 | 0,18 | 6,59±0,23 | 6,47±0,37 | 0,25 | 19,75±5,14 | 19,89±7,07 | 0,95 |
| | Hippocampus | 18 | 15 | 48,17±10,94 | 45,73±8,79 | 0,48 | 9/9 | 6/9 | 0,73 | 7,37±0,64 | 6,54±0,52 | **$3.10^{-4}$** | 6,61±0,21 | 6,43±0,31 | 0,07 | 19,39±5,2 | 19,4±7,2 | 0,81 |
| **GSE62191** | Frontal cortex | 30 | 29 | 44,43±8,57 | 42,17±8,95 | 0,32 | 7/23 | 6/23 | 0,76 | NC/NC | NC/NC | - | 6,45±0,25 | 6,61±0,28 | **0,02** | 31,4±16,93 | 29,97±12,91 | 0,72 |
| **GSE87610** | DLPFC | 19 | 19 | 47,8±10,4 | 45,1 ±8,5 | 0,39 | 9/10 | 9/10 | 1 | 8±0,6 | 7,9±0,7 | 0,64 | 6,6±0,2 | 6,6±0,3 | 1,00 | 19,3±5,3 | 20,1±6,9 | 0,69 |
| **GSE93577** | DLPFC | 19 | 19 | 47,8±10,4 | 45,1±8,5 | 0,39 | 9/10 | 9/10 | 1 | 8±0,6 | 7,9±0,7 | 0,64 | 6,6±0,2 | 6,6±0,3 | 1,00 | 19,3±5,3 | 20,1±6,9 | 0,69 |
| **GSE93987** | DLPFC | 106 | 102 | 48,06±13 | 46,86±12,4 | 0,51 | NC/NC | NC/NC | - | 8,32±0,6 | 8,23±0,6 | 0,28 | 6,76±0,2 | 6,63±0,4 | **$3,7.10^{-3}$** | 17,63±6,1 | 17,98±8,8 | 0,74 |

SZ individuals, *individuals with schizophrenia*; RIN, *RNA integrity number*; PMI, *postmortem interval*; BA, *Brodmann area*; DLPFC, *dorsolateral prefrontal cortex*

**Supplementary Table S3.** Demographic and quality characteristics of the peripheral blood samples

| Study | Brain Region | n subjects Healthy controls | n subjects SZ individuals | Age (years, mean ± SD) | | | Gender (F/M) | | | RIN (mean ± SD) | | | pH (mean ± SD) | | | PMI (hours, mean ± SD) | | |
|---|---|---|---|---|---|---|---|---|---|---|---|---|---|---|---|---|---|---|
| | | | | Healthy controls | SZ individuals | p | Healthy controls | SZ individuals | p | Healthy controls | SZ individuals | p | Healthy controls | SZ individuals | p | Healthy controls | SZ individuals | p |
| **GSE18312** | Blood | 8 | 13 | 45±7 | 43,66±9 | 0,71 | 3/5 | 4/9 | 1 | NC/NC | NC/NC | - | NC/NC | NC/NC | - | NC/NC | NC/NC | - |
| **GSE27383** | Peripheral Blood Mononuclear Cells (PBMC) | 29 | 43 | 23,90±4,08 | 23,02±4,03 | 0,37 | 0/29 | 0/43 | 1 | NC/NC | NC/NC | - | NC/NC | NC/NC | - | NC/NC | NC/NC | - |
| **GSE38485** | Whole blood | 96 | 106 | 39,31±14,19 | 39,58±10,73 | 0,88 | 54/42 | 30/76 | **$6.10^{-5}$** | NC/NC | NC/NC | - | NC/NC | NC/NC | - | NC/NC | NC/NC | - |

SZ individuals, *individuals with schizophrenia*; RIN, *RNA integrity number*; PMI, *postmortem interval*

**Supplementary Table S4.** Results of the differential expression meta-analysis in brain tissues

| **Candidate gene** | ***CSMD1*** | ***CSMD2*** |
|---|---|---|
| **Number of studies combined** | 14 | 13 |
| **Number of observations** | 1054 | 997 |
| **Number of observations derived from HC/SZ individuals** | 536/518 | 507/490 |
| **Effect size (ES)** | 0.07 | 0.22 |
| **95% IC** | [-0.06; 0.20] | [0.05; 0.39] |
| **p value** | 0.31 | **0.013** |
| **FDR-adjusted p value** | **0.31** | **0.026** |
| **p value (Egger test)** | 0.19 | 0.75 |
| **FDR-adjusted p value (Egger test)** | 0.38 | 0.75 |
| **Q test** | 10.7 | 20.3 |
| **p value (Q test)** | 0.64 | 0.062 |
| **FDR-adjusted p value (Q test)** | 0.64 | 0.12 |
| **I2** | 0.00% | 40.8% |

HC. *healthy controls*; SZ individuals. *individuals with schizophrenia*

**Supplementary Table S5.** QM statistics from univariate meta-regression analysis of moderators (age, pH, PMI, RIN) performed for genes differentially expressed in the sex-combined meta-analysis

| Gene | Moderator | QM | p-value |
|---|---|---|---|
| **CSMD1** | | | |
| | pH | 2.7118 | 0.10 |
| | age | 0.0151 | 0.90 |
| | PMI | 0.7944 | 0.37 |
| | RIN | 1.0648 | 0.30 |
| **CSMD2** | | | |
| | pH | 2.6795 | 0.10 |
| | age | 0.0007 | 0.98 |
| | PMI | 0.8162 | 0.37 |
| | RIN | 1.0648 | 0.30 |

PMI, *postmortem interval*; RIN, *RNA integrity number*

**Supplementary Table S6.** Results of the differential expression meta-analysis in the brain tissues of female subjects

| **Candidate gene** | ***CSMD1*** | ***CSMD2*** |
|---|---|---|
| **Number of studies combined** | 10 | 10 |
| **Number of observations** | 189 | 189 |
| **Number of observations derived from HC/SZ individuals** | 101/88 | 101/88 |
| **Effect size (ES)** | 0.21 | 0.31 |
| **95% IC** | [-0.08; 0.50] | [0.02; 0.60] |
| **p value** | 0.15 | **0.037** |
| **FDR-adjusted p value** | 0.15 | 0.074 |
| **p value (Egger test)** | 0.12 | 0.26 |
| **FDR-adjusted p value (Egger test)** | 0.25 | 0.26 |
| **Q test** | 1.53 | 5.89 |
| **p value (Q test)** | 0.99 | 0.75 |
| **FDR-adjusted p value (Q test)** | 0.99 | 0.99 |
| **I2** | 0.0% | 0.0% |

HC. *healthy controls*; SZ individuals. *individuals with schizophrenia*

**Supplementary Table S7.** Results of the differential expression meta-analysis in the brain tissues of male subjects

| **Candidate gene** | ***CSMD1*** | ***CSMD2*** |
|---|---|---|
| **Number of studies combined** | 10 | 10 |
| **Number of observations** | 319 | 321 |
| **Number of observations derived from HC/SZ individuals** | 155/164 | 156/165 |
| **Effect size (ES)** | 0.18 | 0.12 |
| **95% IC** | [-0.04; 0.40] | [-0.16; 0.41] |
| **p value** | 0.12 | 0.39 |
| **FDR-adjusted p value** | 0.23 | 0.39 |
| **p value (Egger test)** | 0.92 | 0.68 |
| **FDR-adjusted p value (Egger test)** | 0.92 | 0.92 |
| **Q test** | 5.68 | 13.61 |
| **p value (Q test)** | 0.77 | 0.14 |
| **FDR-adjusted p value (Q test)** | 0.77 | 0.27 |
| **I2** | 0.0% | 33.9% |

HC. *healthy controls*; SZ individuals. *individuals with schizophrenia*

**Supplementary Table S8.** Results of the differential expression meta-analysis in peripheral blood

| **Candidate gene** | ***CSMD1*** | ***CSMD2*** |
|---|---|---|
| **Number of studies combined** | 3 | 3 |
| **Number of observations** | 288 | 288 |
| **Number of observations derived from HC/SZ individuals** | 133/155 | 133/155 |
| **Effect size (ES)** | -0.19 | -0.08 |
| **95% IC** | [-0.59; 0.20] | [-0.31; 0.16] |
| **p value** | 0.33 | 0.52 |
| **FDR-adjusted p value** | 0.52 | 0.52 |
| **p value (Egger test)** | 0.33 | 0.63 |
| **FDR-adjusted p value (Egger test)** | 0.63 | 0.63 |
| **Q test** | 3.58 | 1.94 |
| **p value (Q test)** | 0.16 | 0.38 |
| **FDR-adjusted p value (Q test)** | 0.33 | 0.38 |
| **I2** | 44.1% | 0.0% |

HC. *healthy controls*; SZ individuals. *individuals with schizophrenia*
